# Population-based mechanistic modeling allows for quantitative predictions of drug responses across cell types

**DOI:** 10.1101/176321

**Authors:** Jingqi QX Gong, Eric A Sobie

**Author notes:** Corresponding author: Eric A. Sobie, Icahn School of Medicine at Mount Sinai, One Gustave Levy Place, Box 1215, New York, NY 10029, USA.

## Abstract

Quantitative mismatches between human physiology and experimental models can present serious limitations for the development of effective therapeutics. We addressed this issue, in the context of cardiac electrophysiology, through mechanistic mathematical modeling combined with statistical analyses. Physiological metrics were simulated in heterogeneous populations describing cardiac myocytes from adult ventricles and those derived from induced pluripotent stem cells (iPSC-CMs). These simulated measures were used to construct a cross-cell type regression model that predicts adult myocyte drug responses from iPSC-CM behaviors. We found that quantitatively accurate predictions of responses to selective or non-selective drugs could be generated based on iPSC-CM responses and that the method can be extended to predict drug responses in diseased as well as healthy cells. This cross-cell type model can be of great value in drug development, and the approach, which can be applied to other fields, represents an important strategy for overcoming experimental model limitations.

## INTRODUCTION

While the goal of much biomedical research is to understand human physiology and pathophysiology, direct human experiments are often infeasible and/or unethical. Because of this, experimental models of human physiology are often required. These can include cell culture models and animal models of human disease. To the extent that they provide a reasonable representation of the human system of interest, the experimental models are useful. However, when a behavior or physiological response in the experimental model does not match the behavior or response seen in the target system, the limitations of the experimental model become a concern. These differences may be qualitative; for instance a drug that is efficacious in a mouse model of disease may fail completely at treating the human disease because mice and humans express different isoforms of a protein. Often, however, these differences are quantitative. For example, a diabetes drug may lower blood glucose in both a mouse model and in diabetic patients, but to different extents.

A strategy for addressing this issue is to build mathematical frameworks that correct for the limitations of the experimental model. In some cases this is trivial -- for instance, the appropriate dose of a drug can often easily be adjusted by a patient’s weight. In other cases empirical correction factors can be derived through painstaking trial and error. However, such correction factors generally only apply under specific conditions, and no general method exists to quantitatively correct for the inaccuracies of how an experimental model will respond to a variety of relevant perturbations.

An example of considerable immediate importance concerns cardiac electrical activity in myocytes derived from induced pluripotent stem cells (iPSC-CMs). Because these cells are a readily-obtainable and renewable source of human cardiac myocytes, they are gaining popularity as a platform to screen drugs for toxicity testing (***Doherty et al., 2015, Gibson et al., 2014, Qu & Vargas, 2015***). However, the cells exhibit immature physiology compared with ventricular myocytes from adult hearts (***Li et al., 2013, van den Heuvel et al., 2014***), and it remains unclear how well drug tests performed in iPSC-CMs will recapitulate the effects observed in human hearts.

We hypothesized that mechanistic, population-based simulations (***Britton et al., 2013, Gemmell et al., 2014, Maltsev & Lakatta, 2013, Marder, 2011, Marder & Taylor, 2011, Sarkar et al., 2012, Walmsley et al., 2013***) could be used to quantitatively map physiological responses between cell types. To test whether this idea was feasible, we attempted to translate drug responses from iPSC-CMs to human adult ventricular myocytes. Beginning with mathematical models of two cell types (***O’Hara et al., 2011, Paci et al., 2013***), we combined simulations of heterogeneous populations with multivariable regression approaches. The resulting model could be used to predict, with quantitative accuracy, drug effects in human adult myocytes based on recordings in iPSC-CMs. Moreover, we found that the approach could be generalized to quantitatively predict effects in diseased myocytes and across multiple species. This strategy is not only practically useful to address contemporary problems in drug development, it provides a framework for addressing the vexing question of experimental model limitations.

## RESULTS

### Human iPSC-CMs and adult myocytes exhibit quantitative differences in their responses to ionic current perturbations

We performed simulations to understand differences between iPSC-CMs and human adult myocytes in the electrophysiological responses to drugs. Fig. 1A and 1B, respectively, show how the two cell types respond to blocking of I_Kr_ and I_CaL_ currents by 25% and 50%. These results demonstrate that the two cell types exhibit differences in baseline action potential (AP) and calcium transient (CaT) morphology, but qualitatively similar responses to ionic current blockade. We quantified the effects of ionic current perturbation by calculating the AP duration at 90% repolarization (APD_90_) and CaT amplitude (CaTA). Figs. 1C and 1D, which plot these outputs over a range of I_Kr_ and I_CaL_ scaling factors, make clear the quantitative divergence between human iPSC-CMs and adult myocytes.

**Figure 1.**
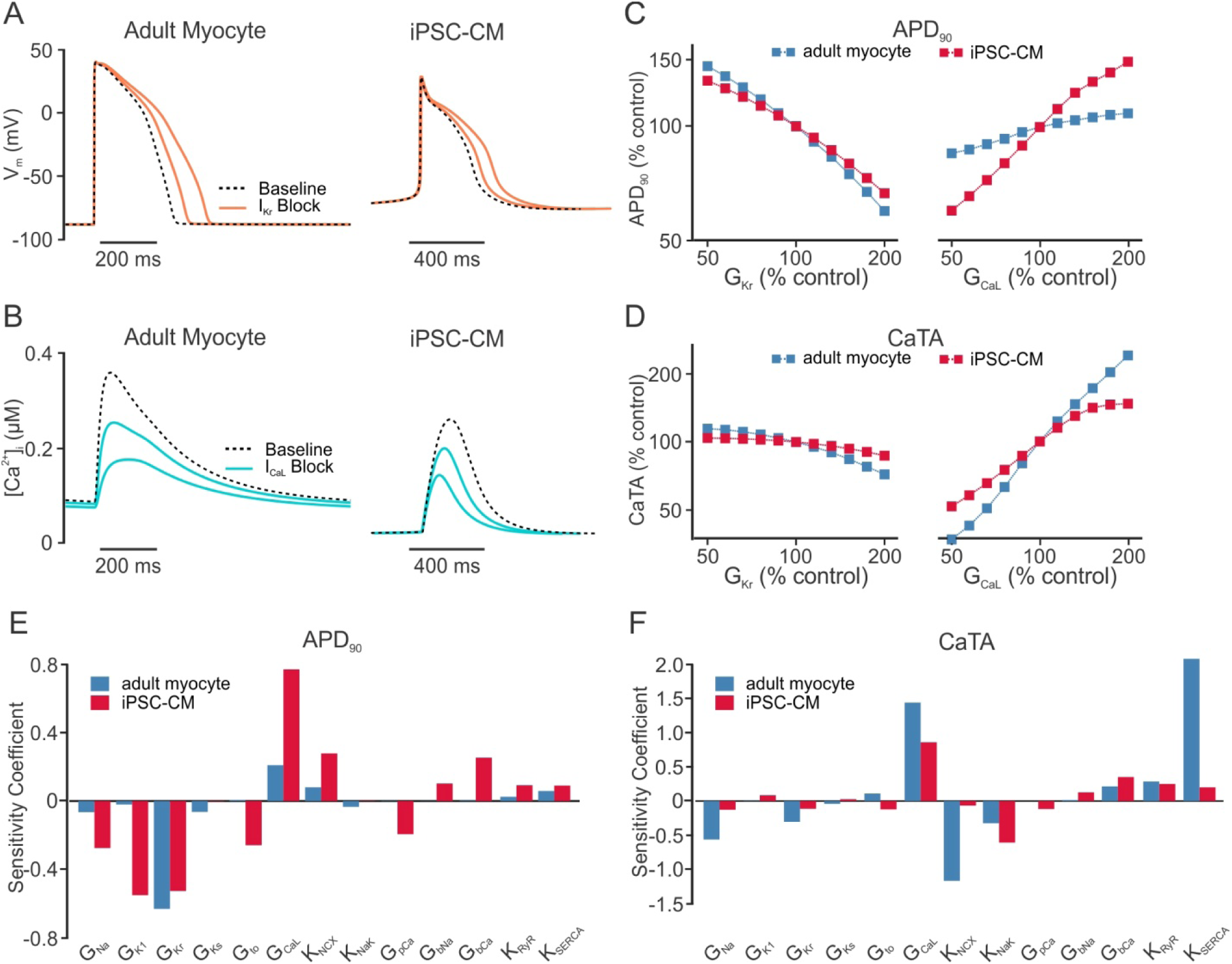
Human adult myocyte and human iPSC-CM responses to perturbations in ion transport pathways. **(A)** Action potential (AP) waveforms simulated in human adult myocyte and iPSC-CM mathematical models before (dashed lines) and after (solid lines) 25% and 50% block of I_Kr_. **(B)** Calcium transient (CaT) time courses of the two cell types at baseline (dashed lines), and after 25% and 50% block of I_CaL_ (solid lines). **(C, D)** Quantification of AP duration at 90% repolarization (APD_90_, **C**) and CaT amplitude (CaTA, **D**) as a function of maximal conductances controlling I_Kr_ (G_Kr_, left) and I_CaL_ (G_CaL_, right) in adult myocyte (blue) and iPSC-CM (red) models. All variables are expressed as a percentage of the control value obtained in the absence of perturbation. **(E, F)** Sensitivity coefficients indicating the extent to which perturbations in each ion transport pathway causes changes in APD_90_ **(E)** and CaTA **(F)**. Coefficients are shown for both adult myocyte (blue) and iPSC-CM (red) models.

To examine how the two cell types respond to a range of ionic current perturbations, we performed parameter sensitivity analyses (***Cummins et al., 2014, Lee et al., 2013, Morotti & Grandi, 2017, Sarkar & Sobie, 2011, Sobie, 2009, Weaver & Wearne, 2008***) of the two models. Across 13 ion channels, pumps, and transporters that are common to both models, we observed marked differences in how perturbations to these ion transport pathways affected APD_90_ and CaTA (Fig. 1E and 1F). These differences highlight that drug responses observed in iPSC-CMs will not necessarily match the responses seen to the same drugs in adult myocytes, and a comprehensive method is therefore required to quantitatively translate physiological responses across cell types.

### A multivariable regression model can translate drug responses across cell types

We developed a statistical model, based on principles of multivariable regression (Fig. 2A), to predict metrics derived from AP and CaT waveforms in adult myocytes from physiological recordings in iPSC-CMs. Heterogeneous *in silico* populations of 600 cells of each type were generated by randomizing the maximal conductance values for 13 ion transport pathways, and Partial Least Squares Regression (PLSR) (***Abdi & Williams, 2013, Geladi & Kowalski, 1986, Kreeger, 2013***) was used to derive a predictive model (see Methods for details). Fig. 2B shows scatter plots of APD_90_ and CaTA obtained with five-fold cross validation. The strong correlation seen between adult model simulation results (abscissa) and regression model predictions (ordinate) indicates that the cross-cell type model is highly accurate (R^2^ = 0.906 for APD_90_; R^2^ = 0.964 for CaTA). Cross validation results of the regression model across 9 additional physiological outputs from AP and CaT are shown in Fig. S1 (Supporting Information). To confirm the model’s practical utility, we simulated 50% blockade of I_Kr_ and I_CaL_ in the baseline iPSC-CM mathematical model, then used the regression model to predict the responses to these perturbations in adult myocytes. The close match in Fig. 2C and 2D between the symbols (regression prediction) and the solid lines (adult model simulation) indicates the accuracy of the model predictions (quantified in Table S3, Supporting Information).

**Figure 2.**
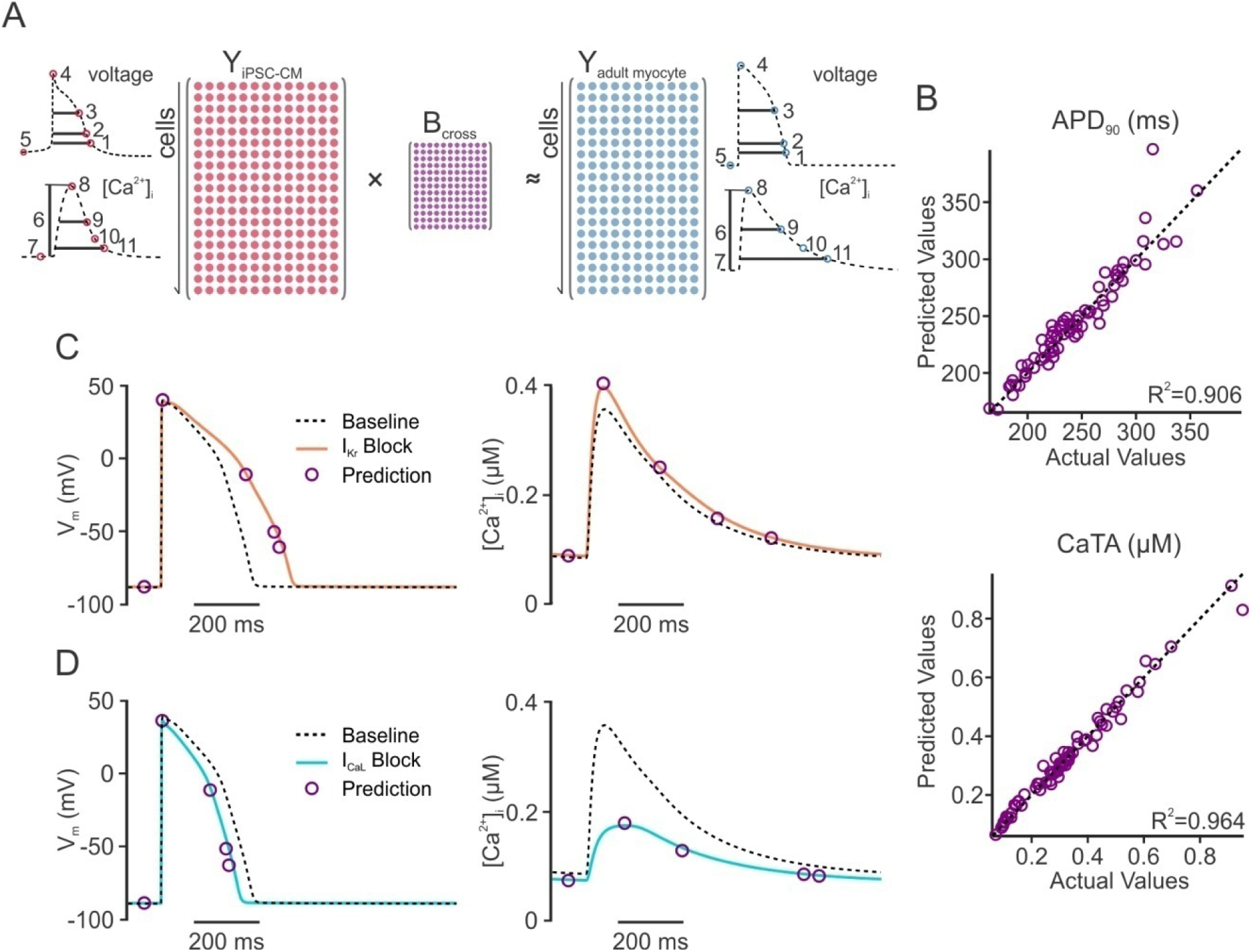
Regression model to predict adult myocyte responses from iPSC-CM physiology. **(A)** Regression strategy for development of a cross-cell type model that maps physiological responses from one cell type (iPSC-CM, left) to another cell type (adult myocyte, right). The resulting regression matrix **B_cross_** serves to generate predictions on adult myocyte responses when measurements are made in iPSC-CM following the same perturbations. Insets: Physiological features quantified from iPSC-CM (left) and adult myocyte (right) simulations. AP features (top): (1) AP duration (APD) at −60 mV; (2) APD at 90% repolarization (APD_90_); (3) APD at 50% repolarization (APD_50_); (4) peak membrane voltage (V_peak_); (5) resting membrane voltage (V_rest_). CaT features (bottom): (6) CaT amplitude (CaTA); (7) resting [Ca^2+^]_I_ (Ca_rest_); (8) peak [Ca^2+^]_I_ (Ca_peak_); (9) CaT duration (CaD) at 50% return to baseline (CaD_50_); (10) CaT decay time; (11) CaT duration (CaD) at 90% return to baseline (CaD_90_). For simulations of iPSC-CM spontaneous (rather than electrically paced) activity, the beating frequency was also quantified and included in the regression model. **(B)** Scatter plots of predictions for adult myocyte APD_90_ (top) and CaTA (bottom), with the actual values from adult myocyte simulations (*abscissa*) versus the cross-cell type predictions (*ordinate*). For clarity, only 100 samples are shown on each of the plots, but the regression was constructed with 600-cell populations, and five-fold cross-validation was performed to calculate R^2^ values. **(C, D)** Adult myocyte AP and CaT responses to 50% block of I_kr_ **(C)** and I_CaL_ **(D)**. Purple circles represent regression model predictions of particular waveform features whereas solid lines indicate numerical simulations.

### Experimental protocols that alter ionic concentrations provide critical information for the regression model

Figure 2 shows results from a regression model in which simulations performed in iPSC-CMs, under 8 experimental conditions (described in Methods), are used to predict responses in adult myocytes. However, some of the 8 protocols may provide partially redundant information, and it is generally unrealistic to record from an individual cell under 8 separate conditions. Therefore, to prioritize efforts and direct experimental design, we evaluated the contribution of each simulated condition to the predictive strength of the cross-cell type model. First we examined iPSC-CM population distributions of APD_90_ (Fig. 3A) and CaTA (Fig. 3B) under different experimental conditions. We observed that, compared with the baseline distributions of spontaneously beating cells (black lines and shaded histograms), some conditions caused minimal changes (e.g. 0.5 Hz pacing, orange histograms) whereas others caused more dramatic shifts in the population distributions (e.g. 2 Hz pacing, blue, and increased extracellular [Na^+^], purple). Based on these results, we hypothesized that conditions which shift these distributions are more informative and contribute more predictive power than those which cause minimal changes.

**Figure 3.**
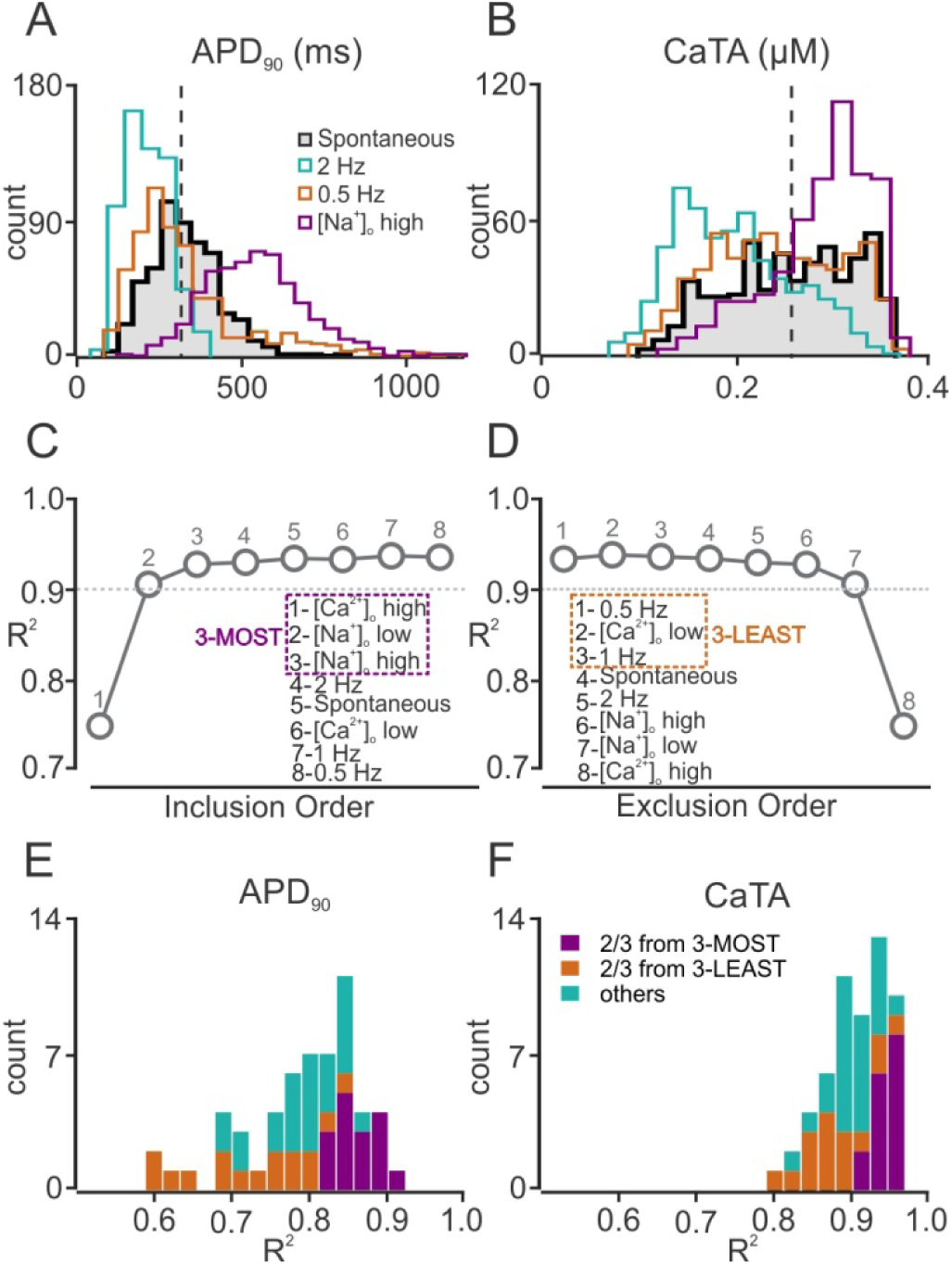
Selection of the most informative iPSC-CM simulation protocols for regression model optimization. **(A, B)** Histograms indicating how APD_90_ **(A)** and CaTA **(B)** vary across a heterogeneous population of iPSC-CMs under different simulated experimental conditions. The black, shaded histogram, representing population behavior with baseline spontaneous contraction is compared with alternative experimental conditions such as 0.5 Hz electrical stimulation (0.5 Hz, orange), 2 Hz electrical stimulation (2.0 Hz, green), and increased (300 mM) extracellular [Na^+^] ([Na^+^]_o_ high, purple). **(C, D)** Averaged R^2^ values across all predicted features with five-fold cross validation, with different numbers of experimental conditions for sequential inclusion **(C)** and sequential exclusion **(D)** methods. These procedures identified the 3 most informative protocols (3-MOST, purple dash square, left) and the 3 least informative protocols (3-LEAST, orange dash square, right). **(E, F)** Distributions of adjusted R^2^ values for APD_90_ **(E)** and CaTA **(F)** of the 56 regression models that can be built by randomly choosing 3 protocols from the initial set of 8. Regression models that select 2 or more protocols from the 3-MOST list (purple) exhibit better predictive power than models that select 2 or more protocols from the 3-LEAST list (orange).

To test this hypothesis and identify the most informative simulation protocols, we constructed regression models based on sequential inclusion (Fig. 3C) or exclusion (Fig. 3D) of additional simulation conditions. With these two complementary approaches, we either included the protocol that led to the greatest improvement in R^2^, or excluded the protocol that caused the smallest decrease in R^2^. The two evaluation approaches led to similar results, and from these we concluded that the 3 most informative simulation conditions (3-MOST protocols) were: 1) an increase in extracellular [Ca^2+^] ([Ca^2+^]_o_ high); 2) a decrease in extracellular [Na^+^] ([Na^+^]_o_ low); and 3) an increase in extracellular [Na^+^] ([Na^+^]_o_ high). Conversely, the 3 least informative protocols (3-LEAST) were: 1) 0.5 Hz pacing, 2) 1 Hz pacing, and 3) a decrease in extracellular [Ca^2+^] ([Ca^2+^]_o_ low). To further validate that the 3-MOST protocols provide complementary information and improve prediction accuracy, we generated all 56 possible regression models that include 3 simulation protocols chosen from the 8, and evaluated each model based on the adjusted R^2^ values for APD_90_ (Fig. 3E) and CaTA (Fig. 3F). Confirming the results of the inclusion and exclusion approaches, we found that combinations with 2 or more protocols from the 3-MOST category (purple bars) achieved higher R^2^ values, compared with models that included 2 or more protocols from 3-LEAST (orange bars), or mixed combinations (blue bars).

Based on these results, our optimized model used in subsequent simulations was built from the three most informative experimental protocols, in addition to spontaneous beating and 2 Hz pacing. The optimized cross-cell type model achieved R^2^=0.903 for APD_90_ and R^2^=0.967 for CaTA following five-fold cross validation.

### Cross-cell type predictions of ionic current blockade are accurate for both selective and nonselective drugs

We next set out to test the ability of the optimized cross-cell type regression model to predict how adult myocytes respond to additional ionic perturbations. These simulations were performed in heterogeneous populations of iPSC-CMs and adult myocytes (100 cells in each group), which allowed us to account for variability between myocytes and to estimate the precision of the predictions. Figs. 4A-D show the simulated effects of I_Kr_ and I_CaL_-blocking drugs over a range of concentrations corresponding to 5-55% channel block. Whether examining the change in APD_90_ (panels A and C) or the change in CaTA (panels B and D), the regression model prediction (purple) provides a much better estimate of the adult myocyte response (gray) than does the change in APD_90_ or CaTA observed directly from iPSC-CMs (cyan). Figs. 4E and 4F show predicted effects of drugs that selectively inhibit 10 ion transport pathways (all simulated at 50% block). With all drugs that cause substantial effects in adult myocytes, the cross cell type predictions (purple) represent a much better prediction of adult myocyte effects (gray) than the straightforward iPSC-CM recordings (cyan).

**Figure 4.**
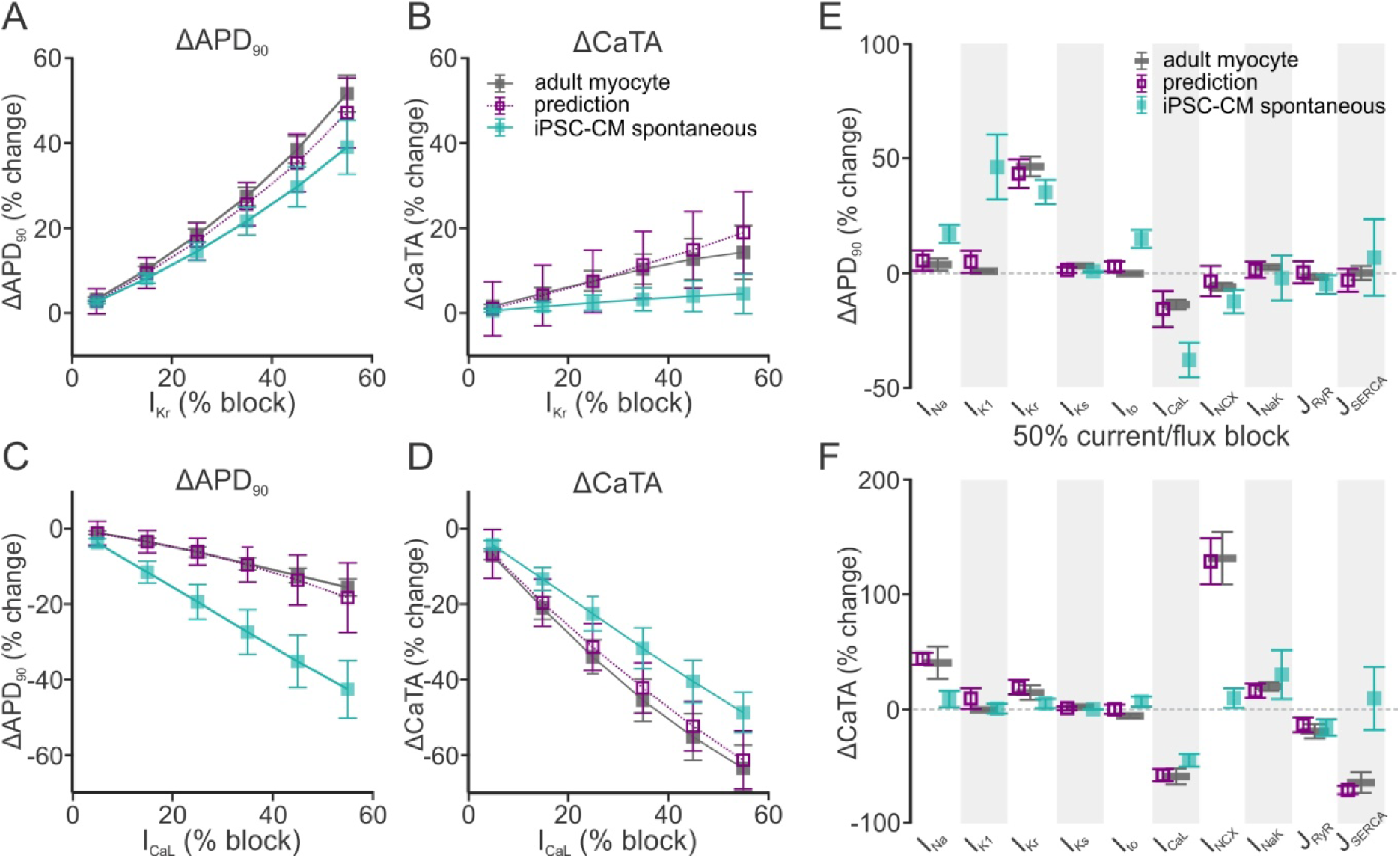
Regression model predictions of how selective ion channel blockers affect adult myocytes. **(A-D)** Simulated selective block of I_Kr_ **(A, B)** and I_CaL_ **(C, D)** varying from 5-55% channel blockade. Responses of APD_90_ **(A, C)** and CaTA **(B, D)** are shown. In each panel, the simulated iPSC-CM response with baseline spontaneous contraction is shown in cyan, the simulated adult myocyte response is shown in dark gray, and the regression model prediction is shown in purple. In all cases, simulations were performed with heterogeneous populations of 100 cells; symbols and error bars represent mean and standard deviation, respectively. **(E, F)** Simulated selective block (50%) of 10 ion transport pathways, with colors as described for panels A-D. Simulated drug-induced changes are shown as percent changes in APD_90_ **(E)** and CaTA **(F)**.

Simulations were also performed to predict the effects of nonselective drugs that inhibited multiple ion transport pathways. For the 10 important ion transport pathways that were simulated (i.e. those in Fig. 4E), we posited 90 hypothetical drugs that blocked 2 of these pathways with different affinities. We assumed that the IC_50_ values for the primary and secondary targets differed by a factor of *e* (2.718), and we simulated drug effects at the lower IC_50_ such that the primary target was inhibited by 50% and the secondary target was inhibited by 27%. Fig. 5 plots, on the abscissa, the percent change simulated in adult myocytes of APD_90_ (Fig. 5, left) and CaTA (Fig. 5, right). The ordinate represents the calculated estimates of these values, taken either directly from spontaneously beating iPSC-CMs (cyan symbols) or the cross-cell type regression model (purple symbols). Simulations were performed in heterogeneous populations of 100 myocytes each, and mean values and population standard deviations are shown. Since, with a perfect predictor, all points would lie on the line of identity, the cross-cell type model clearly produces more accurate predictions of APD_90_ and CaTA than the iPSC-CM recordings. Thus, the cross-cell type regression model accurately predicts the response of adult myocytes to both selective and nonselective drugs.

**Figure 5.**
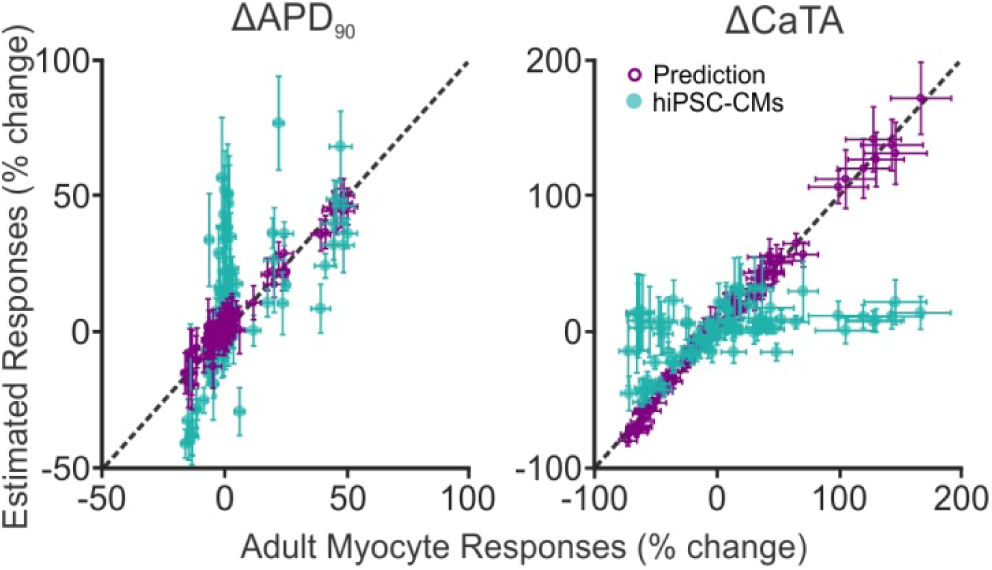
Responses to 90 hypothetical non-selective ion channel blockers. Simulations were performed in adult myocyte and iPSC-CM models of 90 hypothetical drugs that affected two ion transport pathways with different potencies. All simulations were performed in heterogeneous populations of 100 cells each, and mean ± standard deviation is shown. For drug effects on APD_90_ (left) and CaTA (right), simulated adult myocyte responses (*abscissa*) are plotted versus estimated responses (*ordinate*) either directly from iPSC-CM simulations under spontaneous contraction (cyan filled symbols) or from cross-cell type regression model predictions (purple empty symbols). Coefficient of determination (R^2^) was calculated to demonstrate the predictive accuracy. For cross cell type predictions, R^2^=0.957 and 0.987 for APD_90_ and CaTA, respectively. For iPSC-CM spontaneous responses, R^2^=-0.630 and 0.223 for APD_90_ and CaTA, respectively.

### Cross-cell type regression approaches are generalizable for multiple cell types

To test whether the cross-cell type approach was generalizable, we built and tested additional cross-cell type regression using mechanistic models describing ventricular myocytes from additional species. Specifically, using models of rabbit (***Shannon et al., 2004***) and guinea pig (***Livshitz & Rudy, 2009***) ventricular myocytes, we developed regression models that predict: 1) human adult myocyte drug responses from guinea pig or rabbit ventricular physiology; and 2) guinea pig myocyte drug responses from rabbit myocyte physiology, and vice-versa. The overall predictive strength of these models is shown in Supplemental Fig. S3. Validations performed using 50% block of individual ion transport pathways are shown in Fig. 6. Figs. 6A and 6B show how responses to perturbations in the human adult myocytes can be calculated from iPSC-CMs (purple symbols), guinea pig myocytes (blue symbols) or rabbit myocytes (orange symbols). The 3 ionic current perturbations shown are those that caused the largest changes in adult myocytes to either APD_90_ (Fig. 6A) or CaTA (Fig. 6B). In all cases, the prediction of the cross-cell type regression model (open symbols) better approximates the adult response (black bar) compared with direct measurements performed in the alternative cell type (filled symbols). Figs. 6C-6F show examples illustrating that, when clearly different responses to a perturbation are observed in two cells, cross-cell type regression models can correct for the mismatch. For example, 50% block of NCX causes little change to APs in guinea pig myocytes (Fig. 6C, left) but shortens APs in rabbit myocytes (Fig. 6C, right). A cross-cell type regression model built from guinea pig simulations can predict the effects in rabbit, and vice-versa, as quantified in Fig. 6E. As another example, 50% block of SERCA activity reduced CaTA in guinea pig myocytes (Fig. 6D, left) while causing minimal effects in rabbit myocytes (Fig. 6D, right), effects that are accurately captured by the cross-cell type models (Fig. 6F).

**Figure 6.**
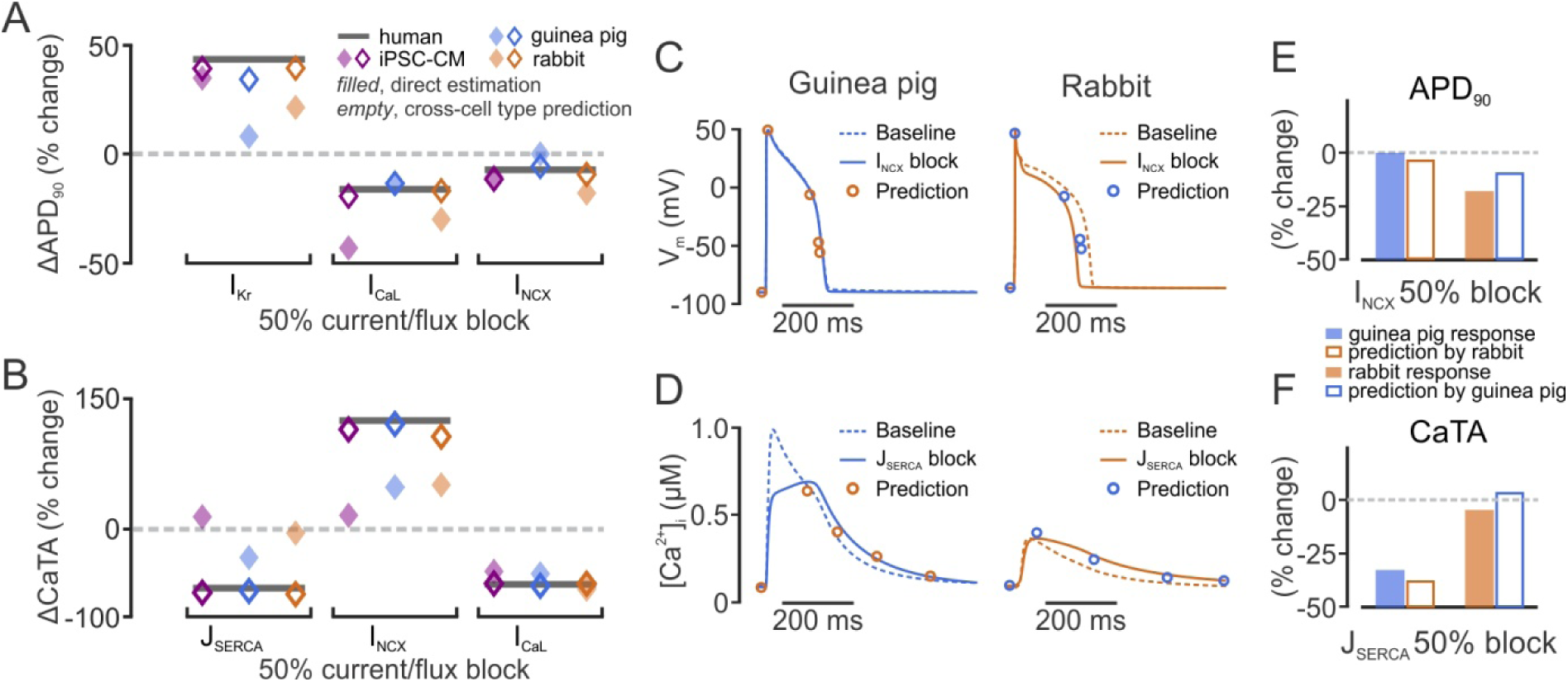
Extension of the cross-cell type regression model concept to additional cell types. **(A, B)** Adult myocyte responses to 50% current/flux block (dark gray bars) were predicted from 3 alternative cell types: iPSC-CM (purple symbols), guinea pig ventricular myocyte (blue symbols), rabbit ventricular myocyte (orange symbols). In each case, filled symbols represent the direct estimate from alternative cell type responses, whereas open symbols represent the cross-cell type regression model predictions. Three ion transport pathways that had large effects in adult myocytes on either APD_90_ **(A)** or CaTA **(B)** are shown. **(C)** Effects of 50% I_NCX_ block on guinea pig and rabbit ventricular action potentials. **(D)** Effects of 50% J_SERCA_ block on guinea pig and rabbit ventricular Ca^2+^ transients. In each case, baseline traces are dashed, perturbed traces are solid, and open circles represent the cross-cell type predictions of waveform features. **(E, F)** Quantification of the results shown in panels C and D.

### Differential drug responses in diseased versus healthy cells can be predicted

In a final step, we determined whether cross-cell type regression could predict differences in drug responses between healthy and diseased cells. Based on published results (***Gomez et al., 2014***), a variant of the O’Hara model was constructed to reproduce molecular and physiological alterations observed in heart failure. Changes in parameters (see ***Gomez et al., 2014*** and Table S5) reflected well-described changes in HF such as reduced SERCA pump activity, upregulation of NCX, and downregulation of I_Ks_. Fig. 7A shows that the HF variant produced slightly longer APs and reduced CaTs, recapitulating the hallmark phenotype of HF myocytes. After generation of a heterogeneous population of HF myocytes (see Methods), PLSR was performed to generate a regression model to predict drug responses in HF myocytes. With this approach, as illustrated in Fig. 7B, recordings obtained under 5 experimental conditions in iPSC-CMs can be used to predict responses in either healthy myocytes (top) or HF myocytes (bottom), using the respective regression matrices for prediction. As an example, these models can successfully predict the differential effects on CaTs seen in healthy and HF cells when NCX is blocked by 40%. This perturbation causes a dramatic increase in CaTA in healthy myocytes (Fig. 7C), but only a small increase in HF myocytes (Fig. 7D), effects that are well predicted by the two regression models (compare symbols with solid lines). Quantification of these effects in Fig. 7E verifies the accuracy of the predictions. Thus, the results shown in Figs. 6 and 7 demonstrate that regression models can accurately translate perturbation effects across cell types, even when the direct effects of a perturbation are dramatically different between the two cell types, including when the question of interest is how different patient populations may respond to the same drug.

**Figure 7.**
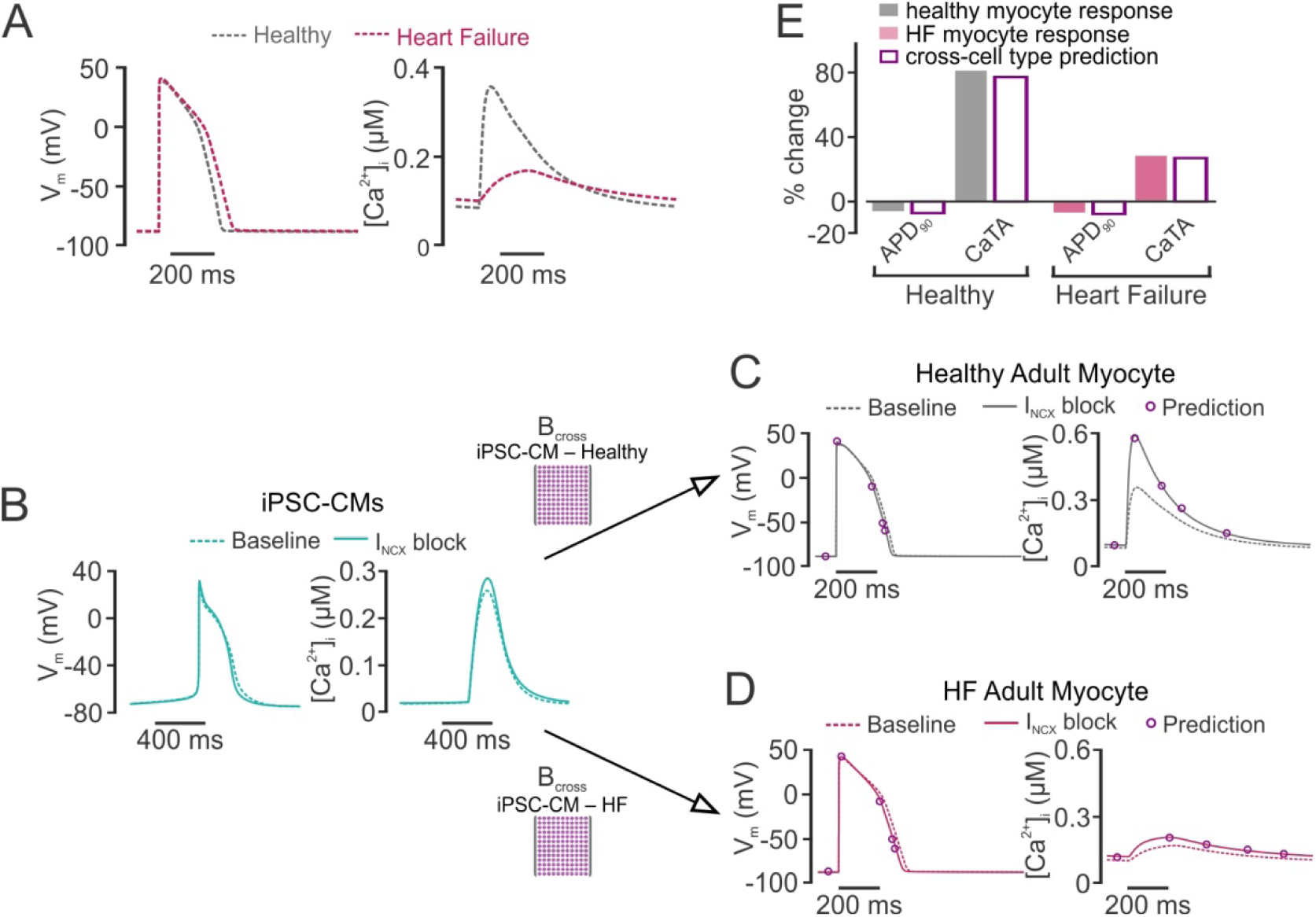
Cross-cell type modeling to predict drug responses in diseased adult myocytes. **(A)** Action potential (AP, left) and Ca^2+^ transient (CaT, right) simulated in the adult myocyte model, with parameters varied to reproduce a heart failure (HF) phenotype, as previously done (***Gomez et al., 2014***). **(B)** Recordings made in iPSC-CMs can be used to predict drug responses in either healthy adult (top) or failing adult (bottom) myocytes, using alternative regression models. **(C)** Regression model accurately predicts that block of I_NCX_ by 40% causes minimal AP shortening and a large increase in CaT amplitude in healthy adult myocytes. **(D)** Regression model accurately predicts that block of I_NCX_ by 40% causes minimal AP shortening and a mild increase in CaT amplitude in failing adult myocytes. **(E)** Quantification of the effects observed in **(C, D)** indicating that cross-cell type model variants can accurately predict drug responses in healthy and diseases populations of adult myocytes. Filled bars represent direct simulations, gray for healthy and pale red for failing. Empty bars are regression model predictions.

## DISCUSSION

In this study we have described a methodology combining mechanistic modeling with statistical analyses to quantitatively translate drug effects across cell types. To develop a predictive model, we first ran simulations with populations of mechanistic cardiac myocyte models. We then performed multivariable regression on the population simulation results -- this regression model allowed us to translate drug responses from iPSC-CMs to human adult myocytes. The model was highly predictive, with cross-validation R^2^ values of 0.906 and 0.964 for APD and CaTA, respectively (Fig. 2B). Moreover, when selective and non-selective blockers of several important ion transport pathways were simulated in iPSC-CMs under multiple conditions, the cross-cell type model predicted adult myocyte responses with quantitative precision. These predictions greatly outperformed a naive approach in which adult myocyte drug responses were assumed to be identical to those seen in iPSC-CMs (Figs. 4 and 5). Importantly, accurate predictions were also obtained when the strategy was applied to ventricular myocyte models from additional species or to a model of diseased human myocytes (Figs. 6 and 7), demonstrating the general utility of the concept.

This novel cross-cell type regression model can address, in a quantitatively rigorous way, differences between an experimental model and the system that is ultimately of interest. The prediction model developed in this study possesses tremendous potential as a practical tool for toxicity testing in drug development, and the overall strategy, which can readily be applied to other fields of research, offers a roadmap for overcoming the limitations that are inherent to experimental models.

### The potential impact of cross-cell type modeling in drug development

Human iPSC-CMs hold considerable promise as a screening platform for assessing how drugs cause either beneficial or deleterious cardiac effects. The application that is most well-developed at present is the use of iPSC-CMs for assessment of drug-induced arrhythmia risk. Specifically, experimental studies with iPSC-CMs are a major pillar of the Comprehensive in vitro Proarrhythmia Assay, or CiPA, a public/private partnership involving the Food and Drug Administration, several pharmaceutical companies, and academic research groups (***Cavero & Holzgrefe, 2014, Colatsky et al., 2016, Fermini et al., 2016, Gintant et al., 2016, Sager et al., 2014***). The goal of CiPA is to replace current pro-arrhythmia tests, which can be non-specific, with a fully in vitro assay involving ionic current measurements, mathematical modeling, and iPSC-CM experiments. A worrisome and unresolved issue within CiPA, however, is the fact that iPSC-CM physiology differs from that of adult myocytes. In general, iPSC-CMs are considered to possess an immature phenotype that resembles fetal heart cells more than adult heart cells (***Li et al., 2013, van den Heuvel et al., 2014***). This is seen both structurally, in the fact that iPSC-CMs lack transverse tubules and a regular organization of sarcomeres (***Bedada et al., 2016***), and functionally, in features such as the contribution of SR Ca^2+^ release to Ca^2+^ transients (***Li et al., 2013***) and the densities of particular K^+^ channels (***Bett et al., 2013***).

Because of these physiological differences, drug responses observed in iPSC-CMs and adult myocytes are also expected to differ (***Ma et al., 2011, Mehta et al., 2013, Paci et al., 2015, Sheng et al., 2012***). The sensitivity analysis shown in Figs. 1E and 1F illustrates these differences by quantifying how changes in any of the shared ion transport pathways influence APD and CaTA in the two model cells. For some effects, such as changes in APD resulting from altered I_Kr_, the two models are largely convergent. For others, however, such as alterations in I_CaL_, dramatic differences are observed. Given that the L-type Ca^2+^ channel is a major ion channel in the CiPA initiative, these discrepancies present a potentially serious problem for the use of iPSC-CMs in drug development. The cross-cell type regression approach described in this study illustrates how these limitations may be overcome. Instead of approximating drug effects directly -- i.e. assuming that drug-induced physiological changes are identical in adult myocytes and iPSC-CMs -- a combination of mechanistic modeling and statistical analyses can be used to develop cross-cell type prediction models, and these in turn can correct for mismatches and provide more accurate predictions.

Of course, another approach to addressing differences between iPSC-CMs and adult myocytes is to improve the iPSC-CM experimental model such that the cell types become more similar. Indeed, recent years have seen many advances to optimize differentiation and produce more mature iPSC-CMs (***Herron et al., 2016, Kadota et al., 2017, Ribeiro et al., 2015***). We note, however, that our strategy is not in competition with these important efforts, instead it is a complementary approach that is likely to become even more urgently needed as maturation methods develop. For one thing, even if a truly optimal iPSC-CM maturation protocol can eventually be identified, cells will almost certainly continue to exhibit some differences with adult cells, and methods to understand these differences quantitatively will still be required. Additionally, the next several years are likely to see a proliferation of alternative methods rather than rapid agreement on a protocol of choice. In this scenario, the field will require ways to translate effects between different iPSC-CM production methods. Finally, a close approximation of the healthy adult ventricular myocyte is not the only cell type that is needed in drug development. Because diseased hearts and healthy hearts may respond differently to the same drugs, we also require methods to understand how therapeutics may affect particular patient populations. Fig. 7 illustrates how, once appropriate regression models have been developed, recordings obtained in a single iPSC-CM preparation can simultaneously predict drug effects in both healthy and failing adult myocytes. Thus, efforts to improve the iPSC-CM experimental model will become more powerful when they are coupled to modeling approaches that can quantitatively synthesize results across different preparations.

### Practical considerations in employing cross-cell type predictions

In addition to developing a model that would be useful for drug development, we also addressed practical considerations required to implement such cross-cell type approach. One of the fundamental principles underlying our model is that studying multiple experimental conditions provides additional information. In other words, simulating only a population of spontaneously contracting iPSC-CMs might not be sufficient to accurately predict drug responses in adult myocytes, but if the same iPSC-CMs are simulated under different pacing protocols, different ionic conditions, etc., then an accurate regression model can be generated. In practical terms, however, it is not always feasible to examine individual cells under a wide range of experimental conditions. Accordingly, we identified the most informative simulation conditions in iPSC-CMs for accurate predictions of adult myocyte responses (Fig. 3). This analysis ranked experimental protocols from the most to the least informative, based on the contribution made by each to the regression model accuracy. An interesting result to arise from this analysis was that altering extracellular ion concentrations appears to be more useful than electrical pacing, a prediction that remains to be tested. In broader terms, however, this shows how the approach we have employed can inform experimental design, by identifying protocols that provide novel information.

In implementing an approach such as we have outlined here, an additional practical consideration that must be addressed is the mathematical model of the iPSC-CM. Although many alternative mathematical models of animal myocytes (***Bondarenko, 2014, Devenyi & Sobie, 2016, Hund & Rudy, 2004***) and human adult ventricular myocytes (***Grandi et al., 2010, ten Tusscher & Panfilov, 2006***) have been published, the Paci et al model (***Paci et al., 2013***) remains, at the present time, the only model of the iPSC-CM. The preferred approach to modeling iPSC-CM physiology will certainly be modified as additional data become available, and as iPSC-CMs developed under particular conditions become more completely characterized. To facilitate the future development of model improvements, an efficient and valuable approach is to choose parameters that match physiological responses (e.g. APs and CaTs) not only under baseline conditions, but in response to perturbations such as augmenting or inhibiting particular ionic currents (***Devenyi et al., 2017, Groenendaal et al., 2015***). Compared with the alternative strategy of characterizing each important ion transport mechanism in every cell type of interest (i.e. fitting models to voltage clamp data), this approach offers the possibility of more rapidly tuning a model to recapitulate experimental data, once the general model structure (i.e. which ion transport mechanisms should be included) is understood (***Krogh-Madsen et al., 2016***). Importantly, these recent studies provide a guide for how to continually improve the cross-cell type approach as additional data are obtained and models of the relevant cell types become more advanced.

### The cross-cell type concept can be extended to additional contexts

Results such as those shown in Figs. 6 and 7 can initially seem quite surprising: if blocking a particular pathway causes minimal effects in one cell type, how can the model successfully reproduce more dramatic effects in another cell type? The answer to this question arises from two important aspects of the cross-cell type regression approach: 1) variability is universally imposed on all the major ion transport pathways when generating the heterogeneous cell populations; and 2) a wide range of simulation results, obtained under multiple experimental conditions, are recorded. These ensure that the underlying statistical associations are robust enough to make accurate predictions, even in cases where obvious differences exist between the source and target cells. Put another way, APD in the adult human myocyte is not simply a function of APD in the iPSC-CM; instead this output depends on an appropriately weighted average of many physiological metrics that can be measured in iPSC-CMs. Our approach therefore advocates for considering all relevant information systematically, and synthesizing based on rigorous statistical analysis, rather than necessarily focusing on particular measurements that are deemed to be most important.

Beyond being a practically useful platform, the cross-cell type regression provides information for future mechanistic studies. Although the extension to rabbit and guinea pig ventricular myocyte models demonstrated that the concept is generalizable (Fig 6), it should be noted that the prediction accuracy was not identical for all models generated. For instance, the worst-performing model, the translation from rabbit to guinea pig myocytes (Fig S3), probably reflects important differences between the two cell types, and these studies can guide future efforts to uncover these mechanistic differences.

More broadly, however, this extension to additional cardiac myocyte models suggests ways that the concept can be further generalized and expanded. The number of cell types seen in the nervous system, for instance, dwarfs what is observed in the heart. However, because different types of neurons employ many of the same channels and receptors to shape their behaviors, an approach such as ours should be feasible for predicting differences. Similarly, the pathways that control processes such as cell division and apoptosis are largely shared between different cell types, although the ways that cells respond to physiological or pharmacological stimuli can be extremely different. For issues such as these, where robust mathematical models already exist (***Albeck et al., 2008, Ciarleglio et al., 2015, Csikasz-Nagy et al., 2006, Drion et al., 2015, Qu et al., 2003, Rinberg et al., 2013, Shin et al., 2014, Sible & Tyson, 2007***), a cross-cell type approach such as we have outlined can be used to predict how behaviors observed in one cell type may or may not translate into a related cell type.

### Study Limitations

Although the results presented in this study are encouraging and provide confidence that the concept can be of practical use, several limitations should be noted. First, in generating the heterogeneous populations, we only varied model parameters controlling ionic current magnitudes, leaving the channel kinetics unaltered. As a result, we could only simulate drug effects through a simple pore block model. Many drugs, however, bind to ion channels in a state-dependent manner, producing effects that our approach could potentially fail to predict. These complexities could be included through the use of more complex Markov models, which have been developed for many critical channels (***Li et al., 2017, Moreno et al., 2011, Silva & Rudy, 2005***). Second, although the regression model performed extremely well with simulated data, the experimental validity of these predictions remains to be tested. Although the ultimate goal of many cardiac experimental models is to predict behaviors in adult human hearts and myocytes, the difficulty of obtaining such tissue for experiments precluded the use of such tests in this study. However, alternative methods for testing the robustness of the concept do exist, for instance by testing the physiological responses to drug treatments across different iPSC-CM preparations. Studies have shown, for instance, that physiological differences exist between iPSC-CMs purchased from alternative vendors (***Blinova et al., 2017, Lu et al., 2017***). These alternative iPSC-CM variants may prove a useful platform for experimentally testing this approach, and the results obtained in such a comparison may help to drive convergence towards a more optimal iPSC-CM experimental model.

### Conclusions

We developed a novel methodology that serves to translate drug effects across cell types with quantitative accuracy, an approach that is likely to be widely useful for addressing differences between experimental models and target systems of interest. Not only is the initial test case we presented, the translation from iPSC-CMs to human adult myocytes, potentially of practical use during toxicity testing to identify pro-arrhythmic drugs, the approach we have outlined can be used to streamline and optimize experimental design. Most important, the methodology we have outlined can easily be applied to other fields in which mechanistic models are well-developed, potentially greatly increasing the impact of the concept and approach.

## ACKNOWLEDGEMENTS

This work was supported by the National Institutes of Health (U01 HL136297 to EAS) and the American Heart Association Predoctoral Fellowship Award (17PRE33670541 to JQXG).

## MATERIALS AND METHODS

### Mathematical models

We developed regression models to predict effects across cell types by using simulations performed with 4 cardiac myocyte models: 1) O’Hara et al. human adult ventricular myocyte model (***O’Hara et al., 2011***); 2) Paci et al. human iPSC-CM model (***Paci et al., 2013***); 3) Livshitz et al. guinea pig ventricular myocyte model (***Livshitz & Rudy, 2009***); and 4) Shannon et al. rabbit ventricular myocyte model (***Shannon et al., 2004***). O’Hara et al. simulations were performed with the model’s endocardial variant, and Paci et al. simulations were performed with the ventricular-like variant. Simulations with the O’Hara et al model were also performed using a model variant in which parameters were altered, based on prior work(***Gomez et al., 2014***), to reproduce molecular and physiological changes observed in heart failure (HF). Parameter modifications made to produce the O’Hara HF variant are shown in Supplementary Table S5.

Each model consists of a system of ordinary differential equations -- models were implemented in MATLAB version R2016b (The MathWorks, Natick, MA), a solver for stiff systems (ode15s) was used for numerical integration, and simulations were performed in a Windows 10 environment.

### Simulations of heterogeneous model populations

To generate heterogeneous populations of models, parameters controlling maximal rates of ion transport were multiplied by scale factors randomly selected from log-normal distributions. Randomly varied model parameters included: fast Na^+^ current (G_Na_), inward rectifier K^+^ current (G_K1_), rapid and slow delayed rectifier K^+^ currents (G_Kr_, G_Ks_), transient outward K^+^ current (G_to_), L-type Ca^2+^ current (G_CaL_), Na^+^-Ca^2+^ exchanger (K_NCX_), Na^+^-K^+^ pump (K_NaK_), sarcolemmal Ca^2+^ pump (G_pCa_), background Na^+^ and Ca^2+^ currents (G_bNa_, G_bCa_), SR Ca^2+^ release flux through ryanodine receptors (K_RyR_), and SR Ca^2+^ uptake via SERCA pumps (K_SERCA_). Baseline values for these model parameters can be found in Supplementary Tables S1-S2. As a result of this scaling, each individual cell in the population has a distinct profile of ion channel/pump/transporter expression levels.

Scale factors were log-normally distributed such that log-transformed values had a mean of zero and a standard deviation of 0.2624. As a result, 95% of the cells in the populations had expression levels ranging between 60-167% of control values. The same set of scale factors was applied to the two cell types used to construct cross-cell type regression models. Simulations were performed for every cell in the population, and features (indicated in Fig. 2, inset) were extracted from the action potential (AP) and calcium transient (CaT) waveforms.

### Simulation protocols

Cells that do not contract spontaneously (adult human, rabbit, and guinea pig myocytes) were allowed to rest for 300 s, then electrically stimulated at 1 Hz for 120 s. The last AP and CaT in the sequence were recorded. For spontaneously contracting iPSC-CMs, the last AP and CaT in a 120 s simulation period were recorded. Additional experimental protocols were simulated in iPSC-CMs, rabbit myocytes, and guinea pig myocytes. These included: 1) electrical pacing at 0.5 Hz and 2 Hz for 120 s; 2) increasing and decreasing extracellular [Ca^2+^] (baseline =1.8 mM, high = 3.0 mM, low = 0.9 mM), 3) increasing and decreasing extracellular [Na^+^] (baseline = 151 mM, high = 300 mM, low = 70 mM); and 4) increasing and decreasing extracellular [K^+^] (baseline = 5.4 mM, high = 10 mM, low = 3 mM). For the protocols that varied extracellular ion concentrations, guinea pig and rabbit myocytes were electrically stimulated whereas spontaneously contracting iPSC-CMs were simulated. We chose these protocols because of the ease with which they can be implemented experimentally.

### Construction and validation of cross-cell type models

Partial Least Squares Regression (PLSR) (***Abdi & Williams, 2013, Geladi & Kowalski, 1986, Kreeger, 2013***) was used to quantitatively relate physiological features in one cell type (the source cell) to those in another cell type (the target cell). To develop these regression models, features quantified from AP and CaT waveforms were placed into matrices. The “input” matrix consisted of features simulated under multiple conditions from the cells in the source population (e.g. iPSC-CM) whereas the “output” matrix consisted of features in the target cells (e.g. adult human myocytes). Each row in these matrices corresponded to a different cell in the population; each column corresponded to a different simulated feature. PLSR was then applied to derive a matrix, **B_cross_**, that translates from one cell type to another and can subsequently be used for predictions. For instance, if the effects of a drug perturbation were recorded in iPSC-CMs, the matrix **B_cross_** could be used to predict the effects of the same drug in adult myocytes.

During model construction, we performed cross-validation to evaluate the prediction accuracy. For example, to perform five-fold cross validation, populations of 600 cells were divided into five subgroups of 120 cells each. Each regression matrix **B_cross_** derived from 4 subgroups (480 cells) was then used to predict behavior in the remaining subgroup. The performance of the cross-cell type model was evaluated using Predicted Residual Sum of Squares, or PRESS (see Supplementary Methods for details).

PLSR is an iterative approach that requires using weighted combinations of input variables (similar to principal components). PLSR is well-suited for datasets in which the columns in the input matrix are correlated; however, it is important to terminate the iterative PLSR algorithm at the appropriate time to avoid overfitting (***Hawkins, 2004, Tetko et al., 1995***). As described in more detail in Supplemental Methods, we chose the number of components of the PLSR model by iteratively determining the smallest number of components that minimized PRESS while simultaneously achieving a high R^2^ value.

### Identification of the most informative and least informative experimental conditions

To determine whether cross-cell type modeling was feasible, we initially constructed a model using simulated results obtained under many experimental conditions. To make this model more suitable for experimental testing, we determined the most informative experimental conditions using sequential inclusion and exclusion methods. First, although we initially simulated 10 experimental conditions (see above), we found that under some conditions, more than 25% of the cells in the population exhibited abnormal dynamics (e.g. afterdepolarizations, failure to repolarize). These were excluded from the determination of the most informative conditions.

For the sequential inclusion method, we generated PLSR models using each of the 8 experimental conditions individually, and selected the one with the highest average R^2^ with 5-fold cross validation. We then added each of the 7 remaining conditions in turn, and selected the one that caused the largest increase in R^2^. This process continued until all 8 experimental conditions had been ranked. Similar criteria were used for the sequential exclusion method, during which we started with all 8 conditions, excluded one condition at a time, then rejected the one that whose exclusion caused the smallest decrease in R^2^ value.

### Simulations of selective and non-selective drugs in heterogeneous populations

We simulated selective and non-selective blockers of ion transport pathways by scaling the model parameters controlling the magnitudes of ionic current or flux. For example, when simulating selective blockade of I_Kr_ by 50% percent, we scaled the conductance G_Kr_ to 50% of its control value. For details of all tested ion transport pathways, see Supplementary Tables S3-S4.

The 90 non-selective hypothetical drugs were designed such that, of the 10 important ion transport pathways (as tested with selective blockers in Fig. 4), each hypothetical drug targets 2 of the 10 pathways with different affinities, and the IC_50_ values for primary and secondary targets differ by a factor of *e* (2.718). To simulate the effects of 90 hypothetical drugs, we scaled the model parameters controlling the maximal magnitude of ion current/flux for primary and secondary targets according to the equation:

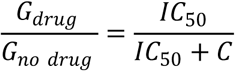

For the 90 hypothetical non-selective blockers, we simulated drug effects at the lower IC_50_, a concentration at which the primary target was inhibited by 50% and the secondary target was inhibited by 27%.

